# SHED-dependent oncogenic signalling of the PEAK3 pseudo-kinase

**DOI:** 10.1101/2021.08.30.457780

**Authors:** Youcef Ounoughene, Elise Fourgous, Yvan Boublik, Estelle Saland, Nathan Guiraud, Christian Recher, Serge Urbach, Philippe Fort, Jean-Emmanuel Sarry, Didier Fesquet, Serge Roche

## Abstract

The PEAK1 and pragmin/PEAK2 pseudo-kinases have emerged as important components of the protein tyrosine kinase pathway implicated in cancer progression. They can signal by a scaffolding mechanism that involves a conserved split helical dimerization (SHED) module. We recently identified PEAK3 as a novel member of this family based on structural homology; however, its signalling mechanism remains unclear. Here, we found that although it can self-associate, PEAK3 shows higher evolutionary divergence than PEAK1/2. Moreover, PEAK3 protein is strongly expressed in human haematopoietic cells, and is upregulated in acute myeloid leukaemia. Functionally, PEAK3 overexpression in U2OS sarcoma cells enhanced their growth and migratory properties, while its silencing in THP1 leukemic cells reduced these effects. Importantly, an intact SHED module was required for these PEAK3 oncogenic activities. Mechanistically, through a phosphokinase survey, we identified PEAK3 as a novel inducer of AKT signalling, independent of growth factor stimulation. Then, proteomic analyses revealed that PEAK3 interacts with the signalling proteins GRB2 and ASAP1/2 and the protein kinase PYK2, and that these interactions require the SHED domain. Moreover, PEAK3 activated PYK2 to promote AKT signalling. Thus, the PEAK1-3 pseudo-kinases may use a conserved SHED-dependent mechanism to activate specific signalling proteins to promote oncogenesis.

## INTRODUCTION

The human kinome includes >50 pseudo-kinases that are predicted to be catalytically inactive due to the lack of important residues required for full enzymatic activity (1). Their mechanistic role in cell signalling remains unclear, but recent structural analyses suggest a scaffolding or allosteric activity by docking additional kinases for efficient protein phosphorylation (1–3). Moreover, some of them have retained active kinase activity through an unconventional mechanism of protein phosphorylation (1–3). These atypical kinases have gained recent interest because they play important roles as active protein kinases in human cancer. For instance, many pseudo-kinases, such as HER3, TRIBL and JAK2 (named JH2), are overexpressed or mutated in various cancer types and contribute to tumour progression (1–3).

The PEAK pseudo-kinases comprise PEAK1, pragmin/PEAK2 and chromosome 19 open reading frame 35 (C19orf35)/PEAK3 (5,6). They include a variable N-terminal sequence with specific interaction motifs (e.g. SH2 and SH3 binding sites) followed by a shared pseudo-kinase domain. Pragmin/PEAK2 (hereafter PEAK2), the family founder, was originally identified as an effector of the small GTPase Rnd2 to mediate cell contraction (7). PEAK1 was identified later as a novel cytoskeletal-associated atypical kinase (8). More recently, using an *in silico* sequence-structure analysis, we identified C19orf35, as an additional family member, hence the PEAK3 name (9). PEAK1/2 are essential components of tyrosine kinase (TK) signalling, leading to cell growth and migration (5,6). Mechanistically, through tyrosine phosphorylation they allow the recruitment of important intracellular signalling effectors, such as GRB2 and SHC (8,10–12). We and other groups have described PEAK1/2 pro-tumour activity (5,6). Specifically, these pseudo-kinases are overexpressed in many epithelial cancer types and contribute to aggressive tumour progression by promoting cancer cell growth and invasion (8,11–18). Importantly, their aberrant expression in tumours may deregulate cancer cell adhesive properties and enable metastatic progression.

Crystallographic studies revealed an important mechanism by which PEAK1/2 pseudokinases may regulate oncogenic pathways (9,19,20). In addition to a classical protein kinase fold harbouring an occluded ATP binding site, three crystal structures revealed an original dimerization domain, named split helical dimerization (SHED) (9,19,20). This conserved module comprises a long helix N-terminal to the pseudo-kinase domain and three additional helices from the C-terminal extension that form an “XL”-shaped helical bundle. The SHED-based dimerization mechanism regulates PEAK1/2 homo- and hetero-dimerization that might influence PEAK signalling specificity (21). However, it is not known whether the SHED domain defines an essential mechanism of PEAK signalling.

PEAK3 was identified as a member of the PEAK family because its peptide sequence predicts a similar degraded ATP-binding site in the TK flanked by the SHED module (9). Moreover, PEAK3 self-associates in a SHED-dependent manner (22), but its biological activity is poorly characterized. Recently, Lopez et al reported that PEAK3 expression can prevent CRKII-dependent membrane ruffling (22), suggesting a negative function in cell migration, unlike PEAK1/2. Here, we investigated PEAK3 functional role in cancer. We found that PEAK3 promotes cancer cell growth and migration. Moreover, the SHED module has an important function in these effects through a unique PEAK3/PYK2/AKT signalling cascade. Our findings support the existence of a common SHED-dependent mechanism of PEAK1-3 oncogenic activity mediated by the recruitment of a unique set of signalling proteins.

## MATERIAL AND METHODS

### Antibodies

The anti-PEAK3 antibody was generated against a bacterially produced recombinant His-TRX-PEAK3 fusion protein (38). Briefly, the His-tagged protein was purified in denaturing condition in the presence of 8M guanidine hydrochloride, followed by extensive dialysis against PBS before rabbit immunization (39). Anti-PEAK3 antibodies were affinity-purified against His-PEAK3 that was blotted on a PVDF membrane (40). Anti-ERK1/2, anti-ERK1/2 pT202/Y204, anti-AKT, anti-AKT pS473, anti-pSer/pThr AKT pan-substrate, anti-GRB2, anti-ASAP1/2, anti-CRK II, anti-PYK2, anti-pPYK2 pTyr402 and anti-PYK2 pTyr881 antibodies were from CST; the anti-actin and anti-tubulin and 4G10 antibody was a gift from N Morin (CRBM); the anti-MYC antibody 9E10 was from Origene, the anti-Strep-tag antibody from Biorad, the anti-rabbit IgG-HRP and anti-mouse IgG-HRP antibodies from GE Healthcare, and the anti-HA antibody 12CA5 from Sigma. The PYK2 inhibitor PF-431396 (1 μM) was from Selleck Biochem, and the pan-PI3K inhibitor LY294002 (10 μM) from Sigma.

### Constructs

The codon-optimized PEAK3 (GenBank NM198532) in the pDONR221 vector was obtained from DNASU (catalogue number HsCD00813413) and subcloned in the pBabe-puro GFP-N-term pCDNA-T0-FRT-Streptag (41) and pSBbi-PUR Sleeping Beauty-based expression vectors (Addgene #6023, a gift from E. Kowarz) (42). The PEAK3 A436E mutant was generated with the QuickChange Site-Directed Mutagenesis Kit (Stratagene). The pCDNA3.1-V5-PTK2B (PYK2) plasmid was a gift from Kai Johnsson (Addgene plasmid # 127233). The siRNAs were: 5’TTCTCCGAACGTGTCACGTTT3’ (siCTRL) and 5’GTATAGCAACCTTGGTCAGAT 3’ (siPEAK3) (31).

### Phylogenetic analyses

Nucleic and protein PEAK sequences were retrieved from the NCBI annotated non-redundant database (http://www.ncbi.nlm.nih.gov) using NCBI PHI-BLAST and also BLAST and the Annotation search tools available in the Geneious 11.1.5 software package (Biomatters, http://www.geneious.com). Accession numbers are listed in Table S1. Protein sequences were aligned using MAAFT v7.450 (43). Protein multiple sequence alignments (MSA) were processed with BMGE (Block Mapping and Gathering with Entropy) (44) and a cut-off value = 0.6. Phylogenetic trees were estimated with PhyML (45), using the General Time Reversible model with invariant and gamma decorations, and MrBayes (46) with two independent runs of four Markov chains for 1,000,000 generations samples every 200 generations. Sample trees from the first 100,000 generations were discarded as “burn-in”. At the end of the two runs, the split frequency standard deviations were ~1.5×10-3 and the effective sample sizes were >2600. The non-synonymous to silent substitution ratios (ω= dN/dS) were calculated using PAML 4.4 (47). Nucleic MSAs were based on protein alignments. The “one-ratio” (i.e. a single ω ratio for the entire tree) and the two “two-ratio” models (i.e. distinct ω values for two or three PEAK branches) were compared using the likelihood-ratio test.

### Data mining

Transcriptomic signatures and data mining from two PEAK3 signatures were generated from the transcriptomes of the acute myeloid leukaemia (AML) patient samples and high versus low PEAK3 expression from TCGA transcriptomic data (gepia.cancer-pku.cn) and BEATAML database (GSE6891) and correlated with well-characterized oncogenic somatic mutations.

### Cell cultures and transfections

Adherent cell lines (ACTT; Rockville, MD) were cultured at 37°C and 5% CO_2_ in a humidified incubator in Dulbecco’s Modified Eagle’s Medium GlutaMAX. Blood cell-derived cell lines were grown in RPMI (Invitrogen) supplemented with 10% foetal calf serum (FCS), 100 U/ml of penicillin and 100 μg/ml of streptomycin. Cell transfections were performed as described in (48). U2OS, HeLa and THP1 cell lines that stably express GFP- and Strep-tagged (ST)-PEAK3 were obtained by transfection of the pBABE-GFP-PEAK3 and pSBbi-PUR-ST-PEAK3 constructs, respectively, with TurboFect^™^, followed by puromycin selection. HEK293T cells were transiently transfected with jetPEI® (Polyplus-transfection) according to the manufacturer’s instructions. For siRNA transfection, 2.10^5^ cells were seeded in 6-well plates and transfected with 20 nmol of siRNAs and 9 μl of Lipofectamine® RNAi Max, according to the manufacturer’s protocol (ThermoFisher Scientific), for 3 days before use.

### Human samples

Primary AML samples from patients (with annotated clinical and biological characteristics, treatment and outcome data) were obtained from the Biobank of the Haematology Clinical Department, Toulouse University Hospital, France, (HIMIP collection, BB-0033-00060; INSERM-U1037) headed by Prof C. Récher. The recording in the database of clinical and biological data have been declared to the CNIL (Comité National Informatique et Libertés; the French Committee for the protection of personal data). Samples are routinely characterized by morphologic, immunophenotypic, cytogenetic and molecular (FLT3-ITD, NPM1, CEBPA, DNMT3A, KIT, IDH1, IDH2 mutations) analyses.

### Biochemistry and phospho-kinase array

Immunoprecipitation and immunoblotting were performed as described in (48). Briefly, cells were lysed at 4°C with lysis buffer (20mM Hepes pH7.5, 150mM NaCl, 0.5% Triton X-100, 6mM β-octylglucoside, 10μg/ml aprotinin, 20μM leupeptin, 1mM NaF, 1mM DTT, and 100μM sodium orthovanadate). Then, 20-50 μg of whole cell lysates were loaded on SDS-PAGE gels and after electrophoretic separation, were transferred onto Immobilon membranes (Millipore). After incubation with the relevant antibodies, detection was performed with the ECL System (Amersham Biosciences).

The Proteome Profiler Human Phospho-Kinase Array Kit was purchased from R&D Systems. THP1 cells were lysed, and 300 μg of protein lysates were used for western blotting according to the manufacturer’s protocol. Signals were quantified with the Amersham Imager 600 (GE Healthcare) from two independent biological replicates.

### Proteomic analyses

Interactomic analyses were performed as described in (49). Briefly, cell lysates were incubated with streptavidin beads (Lifescience) overnight, and washed three times with lysis buffer and once with washing buffer (50mM Tris pH7.5 and 50mM NaCl). PEAK3 complexes were eluted with 2.5mM biotin in 50mM Tris pH8 and 0.5 mM EDTA, and processed for mass spectrometry (MS) analysis. Peptides obtained after digestion were analyzed using a Qexactive-HFX system coupled with a RSLC-U3000 nano HPLC. Samples were desalted and pre-concentrated on-line on a Pepmap® precolumn (0.3 mm x 10 mm). A gradient consisting of 6-25% B for 100 min, 25-40% B for 20 min, 40-90% B for 2 min (A = 0.1% formic acid; B = 0.1 % formic acid in 80% acetonitrile) at 300 nl/min was used to elute peptides from the capillary reverse-phase column (0.075 mm x 250 mm; Pepmap®, ThermoFisherScientific), fitted with a stainless steel emitter (Thermo Sientific). Spectra were acquired with the instrument operating in the data-dependent acquisition mode throughout the HPLC gradient. MS scans were acquired with resolution set at 60,000. Up to twelve of the most intense ions per cycle were fragmented and analyzed using a resolution of 30,0000. Peptide fragmentation was performed using nitrogen gas on the most abundant and at least doubly charged ions detected in the initial MS scan and an dynamic exclusion time of 20s. Analysis was performed with the MaxQuant software (version 1.5.5.1). All MS/MS spectra were searched using Andromeda against a decoy database consisting of a combination of the *Homo sapiens* reference proteome (release 2019_02 www.uniprot.org) and of 250 classical contaminants, containing forward and reverse entities. A maximum of two missed cleavages was allowed. The search was performed with oxidation (M) and acetyl (protein N-term) as variable modifications, and carbamidomethyl (C) as fixed modification. FDR was set at 0.01 for peptides and proteins, and the minimum peptide length was 7. The *t*-test and graphical representation were performed with Perseus (v 1.6.10.43) and standart parameters.

### Cell growth and migration

Cell growth was measured by Sulforhodamide B staining (Sigma Aldrich), and invasion was assessed using Boyden chambers coated with Matrigel (1mg/ml) (12). Cell transmigration was monitored as described in (50).

### Cell imaging

Cells seeded on microscope coverslips were fixed with 4% paraformaldehyde in Triton-X100-BRB80 buffer (0.5% Triton-X100, 80mM PIPES pH 6.8, 1mM MgCl_2_, 1mM EGTA) containing 5% FCS at 37°C for 20min. Cells were incubated with the indicated primary antibodies and the corresponding secondary antibodies labelled with Alexa Fluor dyes (488 or 594nm) and DAPI diluted in PBS buffer. After mounting with ProLong Antidafe Mountant (ThermoFisher), cells were observed with a DMR A microscope and a PL APO 63× oil objective (1.32 NA). Images were captured using a piezzo stepper (E662 LVPTZ amplifier; Servo) and a cooled CCD Micromax camera (1,300 × 1,030 pixels, RS; Princeton Instruments Inc.) driven by MetaMorph (v 4.5; Universal Imaging Corp.). Images were processed with ImageJ (https://imagej.nih.gov/ij/).

### Statistical analysis

All statistical analyses were performed on data from at least three independent experiments with GraphPad Prism. Data are presented as the mean ± standard deviation (SD). When distribution was normal (assessed with the Shapiro-Wilk test), the two-tailed *t* test was used for between-group comparisons. Otherwise, the Mann-Whitney test was used.

## RESULTS

### PEAK3 evolutionary conservation

Unlike the human genome (three PEAK genes), we found a single PEAK gene in the genomes of early metazoan species, from placozoans (*Trichoplax adherens*) to early chordates (the lancelet *Branchiostoma*) (22) (Fig. 1a) (9). To better understand the PEAK family ontology, we searched for homologous sequences in key evolutionary species. We identified two PEAK members in cyclostomes (the lamprey *Petromyzon marinus* and the hagfish *Eptatretus burgeri*), and three members in cartilaginous fishes (e.g. the elephant shark *Callorhinchus milii*), bony fishes (e.g. *Takifugu rubripes*), and the coelacanth *Latimeria chalumnae*. We then extracted the three homologous sequences in additional Tetrapoda species and performed a clustering analysis of MSAs using two probabilistic algorithms (maximum-likelihood and Bayesian). We used the related PTEN-induced kinases (PINK) as external group to root the tree. The clustering tree globally followed the species phylogeny, and showed the presence of a well-supported node leading to three PEAK1-3 clusters in vertebrates (Fig. 1b). The branching of the three nodes suggests that a first duplication led to PEAK1/2 and PEAK3, followed by the PEAK1/2 duplication into PEAK1 and PEAK2. This shows that PEAK1-3 emerged at the onset of vertebrates, and not in Sauropsida as previously proposed (22). The presence of only PEAK1-like proteins in cyclostomes suggests that they lost PEAK2 and PEAK3. However, the lower conservation of PEAK3, as indicated by the longer internal branches, may have blurred the branching of the three groups to such an extent that the exact duplication order can no longer be deduced.

**Figure 1.**
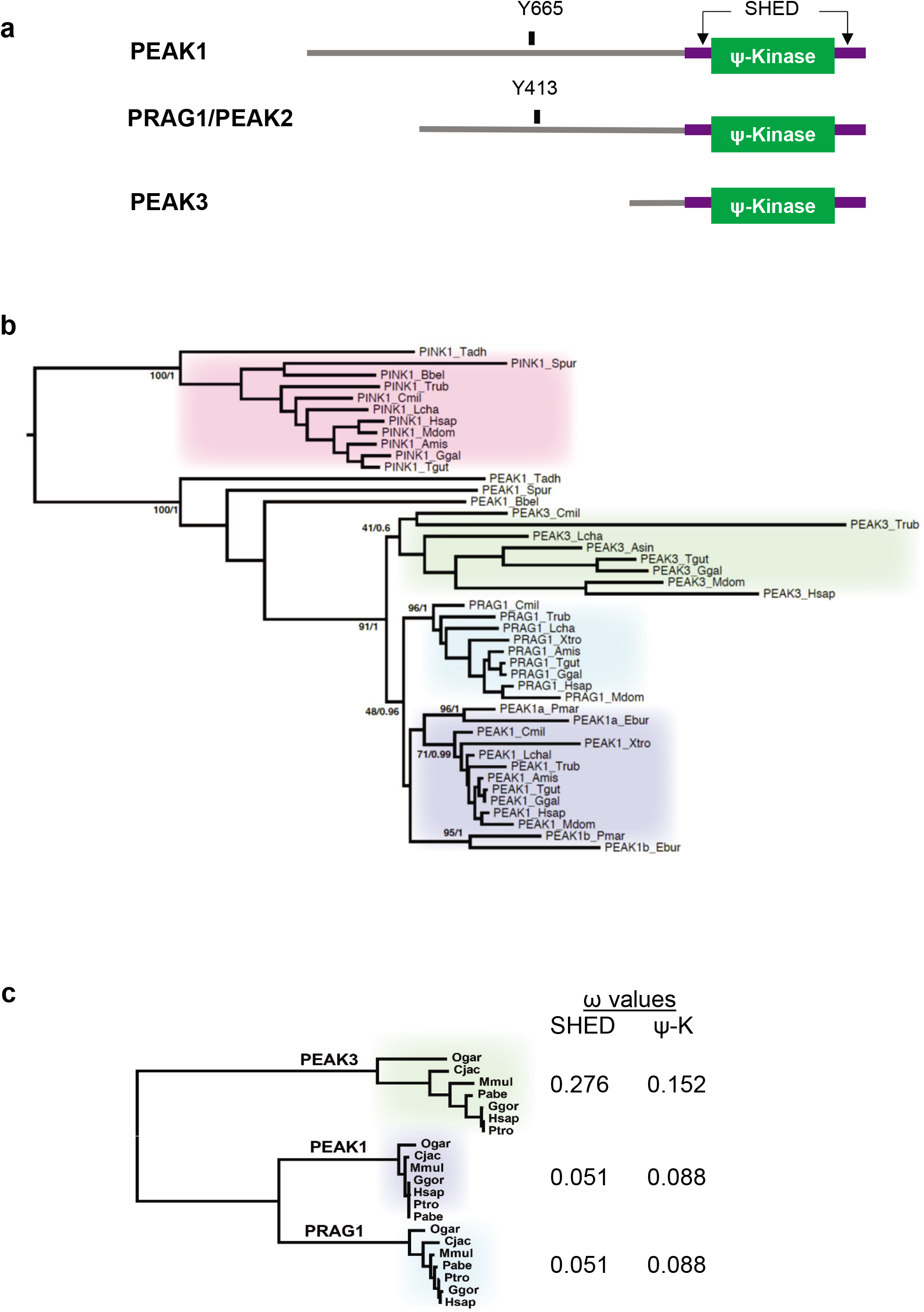
Phylogenetic analysis of PEAK pseudo-kinases. **a**. Modular structure of PEAK1-3 pseudo-kinases. The Split Helical Dimerization (SHED) module (made of αN, αJ, αK and αN helices), the pseudo-kinase (ϕ-Kinase), and the main tyrosine phosphorylation sites are highlighted. **b**. Phylogenetic cladogram of PEAK proteins. The tree was deduced from a multiple amino acid sequence alignment of kinase and pseudo-kinase domains and processed by PhyML and MrBayes analysis. PTEN-induced kinases (PINK) were included as external group to root the tree. Only supports for key phylogenetic nodes are indicated (bootstrap proportion/posterior probability). Species and sequences (accession numbers) used for the alignment are listed in Table S1. **c**. Analysis of the non-synonymous to silent substitution ratios (ω) of the SHED and pseudo-kinase (ΨK) domains in primate PEAKs. The nucleic sequences of the domains were aligned based on translation and processed with codeml to estimate the ω-values. The phylogenetic tree on the left was generated by PhyML analysis of a multiple alignment of the ΨK amino acid sequences. Species and sequences (accession numbers) used for the alignment are listed in Table S1.

To firmly establish the lower conservation of PEAK3 compared with PEAK1/2, we examined the selective constraints exerted on the SHED and pseudo-kinase (ΨK) domains. We calculated the dN/dS ω ratio from a MSA of seven primate species that covered 74 MY of evolution. Neutral selection is associated with ω values close to 1, while values below or above 1 indicate purifying and diversifying selection, respectively. The selection exerted on the SHED and ΨK domains was lower for PEAK3 than for PEAK1 and PEAK2 (Fig. 1c), in agreement with the difference in branch length. The lowest PEAK3 conservation also agrees with the observation that the corresponding gene was lost in several taxa, for instance Myomorpha in rodents (Table S1). The maintenance of duplicated genes is often associated with neo- or sub-functionalization (i.e. the acquisition of new functions or differential tissue expression, respectively) (23).

Then, we investigated *PEAK1-3* mRNA expression using the Human Protein Atlas (24) and found that *PEAK1* was ubiquitously expressed and particularly in spleen; *PEAK2* expression was more restricted, with the highest expression in cerebellum; and *PEAK3* was nearly exclusively expressed in lymphoid tissues (spleen, lymph nodes) and granulocytes (Fig. S1).

Overall, these data indicate that although PEAK3 is much less evolutionary conserved than its paralogues PEAK1 and PEAK2, it underwent purifying selection, indicative of a physiological function, possibly in blood cells.

### PEAK3 overexpression in acute myeloid leukaemia

Next, to investigate PEAK3 protein expression in human cells, we generated a polyclonal antibody against full-length recombinant PEAK3. This antibody could detect in western blotting PEAK3 in lysates from PEAK3-overexpressing U2OS and HeLa cells (Fig. S2a). It also recognized an endogenous protein of about 50KDa in lysates of THP1 (AML) cells (Fig. S2b). We confirmed that this protein was PEAK3 because the 50 KDa band disappeared upon siRNA-mediated PEAK3 knockdown (Fig S2b). We then used this antibody to assess PEAK3 expression in various human cell lines. We did not detect endogenous PEAK3 in the mesenchymal and epithelia cell lines tested (i.e. MCF7, U2OS, HeLa, SW620 and HCT116) (Fig. S2c), but only in haematopoietic cell lines, specifically in AML cell lines (i.e. MV4-11, THP1, MOLM14, and OCI-AML3) (Fig. S2c and Fig. 2a). Consistent with this finding, we observed aberrant PEAK3 protein expression in 40% of the tested AML patient samples (8/20) (Fig. 2b and Table S2). Analysis of *PEAK3* transcript level from TCGA transcriptomic data (gepia.cancer-pku.cn) confirmed that *PEAK3* was upregulated in AML (Fig. 1c), particularly in the M4 and M5 subtypes (Fig. S3a). PEAK3 aberrant expression was preferentially associated with specific oncogenic mutations, i.e. DNTM3A mutations, but not with FLT3-TD mutations (Fig. S3b). Our findings indicate that PEAK3 is expressed in haematopoietic cells and that its expression can be deregulated in AML, suggesting a pro-tumour function.

**Figure 2.**
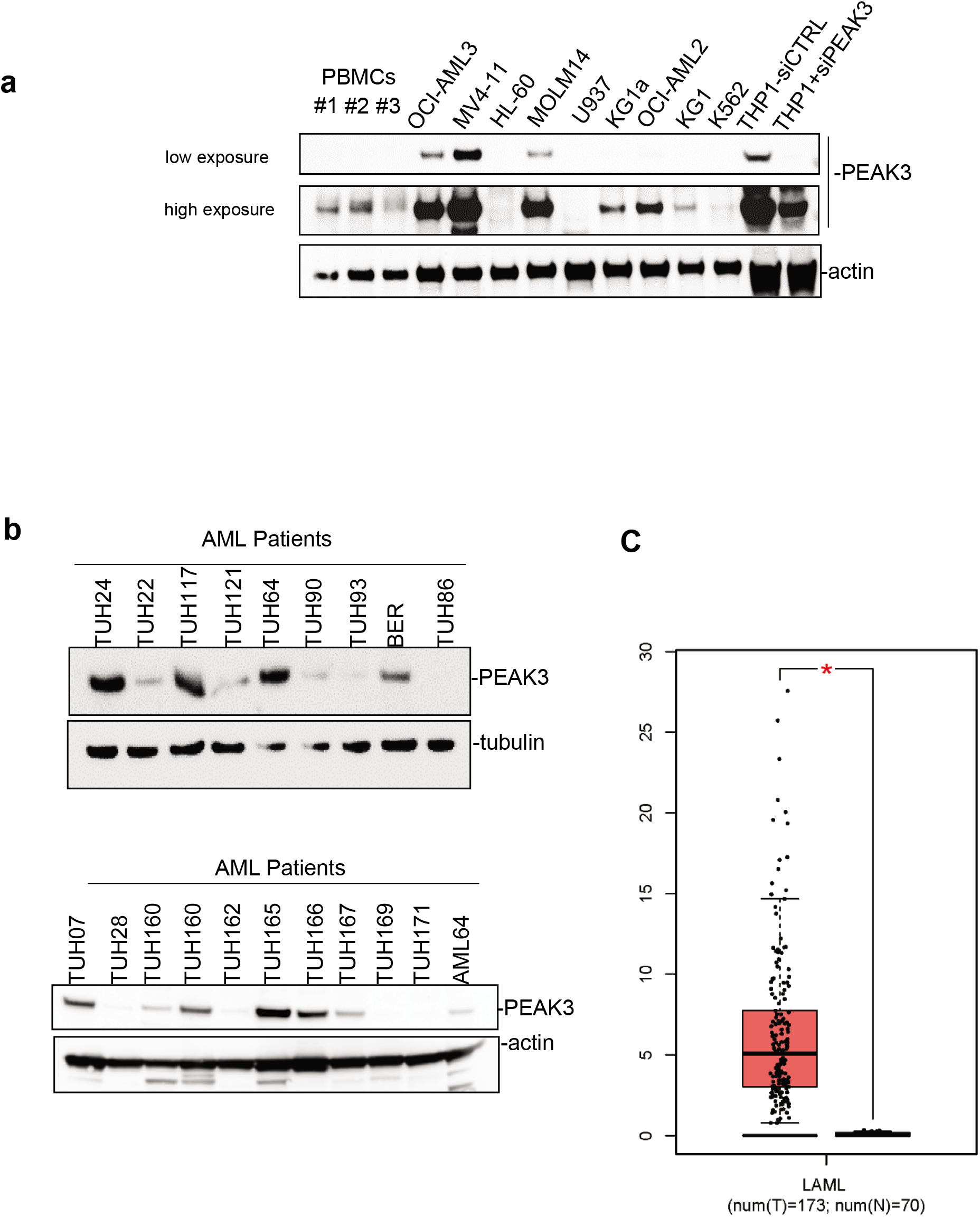
PEAK3 expression in AML. **a**. PEAK3 protein expression in the indicated blood cell lines revealed by western blotting with an anti-PEAK3 antibody. The antibody specificity was confirmed using THP1 cells transfected with a siRNA targeting PEAK3. Both low and longer exposure are shown. PBMCs, peripheral blood mononuclear cells. **b**. PEAK3 expression in AML patient samples. The level of actin or tubulin in also shown. **c.** PEAK3 upregulation in AML patient samples. The panel shows the relative level of PEAK3 transcripts in 173 AML samples (T) and 70 normal samples (N), (gepia.cancer-pku.cn). *p<0.05 (Student’s *t* test).

### SHED-dependent oncogenic activity of PEAK3

We then investigated whether PEAK3 had pro-tumour functions. PEAK3 overexpression in U2OS cells (Fig. 3a) and in THP1 cells (Fig. 3c) increased cell growth and invasion in Matrigel-coated Boyden chambers (Fig. 3b). These effects were abrogated in both cell lines by overexpression of PEAK3 harbouring the A436E mutation in the C-terminal extension of the SHED module (Fig. 3a-c), which destabilizes its dimerization (Fig. 3a and Fig. S4a)(22). Similarly, siRNA-mediated *PEAK3* silencing in THP1 cells reduced their growth and migration, in Boyden chambers coated with an endothelial cell monolayer (Fig 3d). These findings indicate that in human cancer cells, PEAK3 displays oncogenic activity that requires an intact SHED module. As PEAK1/2 localizes at focal adhesions (FAs)(8,14), we investigated PEAK3 localization by direct fluorescence analysis of U2OS cells that stably express GFP-PEAK3 and demonstrated PEAK3 localization at FAs (Fig. S5a), and partial co-localization with paxillin (Fig. S5a). We confirmed PEAK3 localization at FAs by indirect immunofluorescence using an anti-HA antibody in U2OS cells that stably express HA-ST-PEAK3 (Fig. S5b). Additionally, both imaging methods showed a strong PEAK3 nuclear localization (Fig. S5). Conversely, the PEAK3 A436E mutant did not display FA localization, but was still localized in the nucleus (Fig S5a-b). These observations indicate that PEAK3 localizes at FAs, like PEAK1/2, and that this requires its SHED dimeric module.

**Figure 3.**
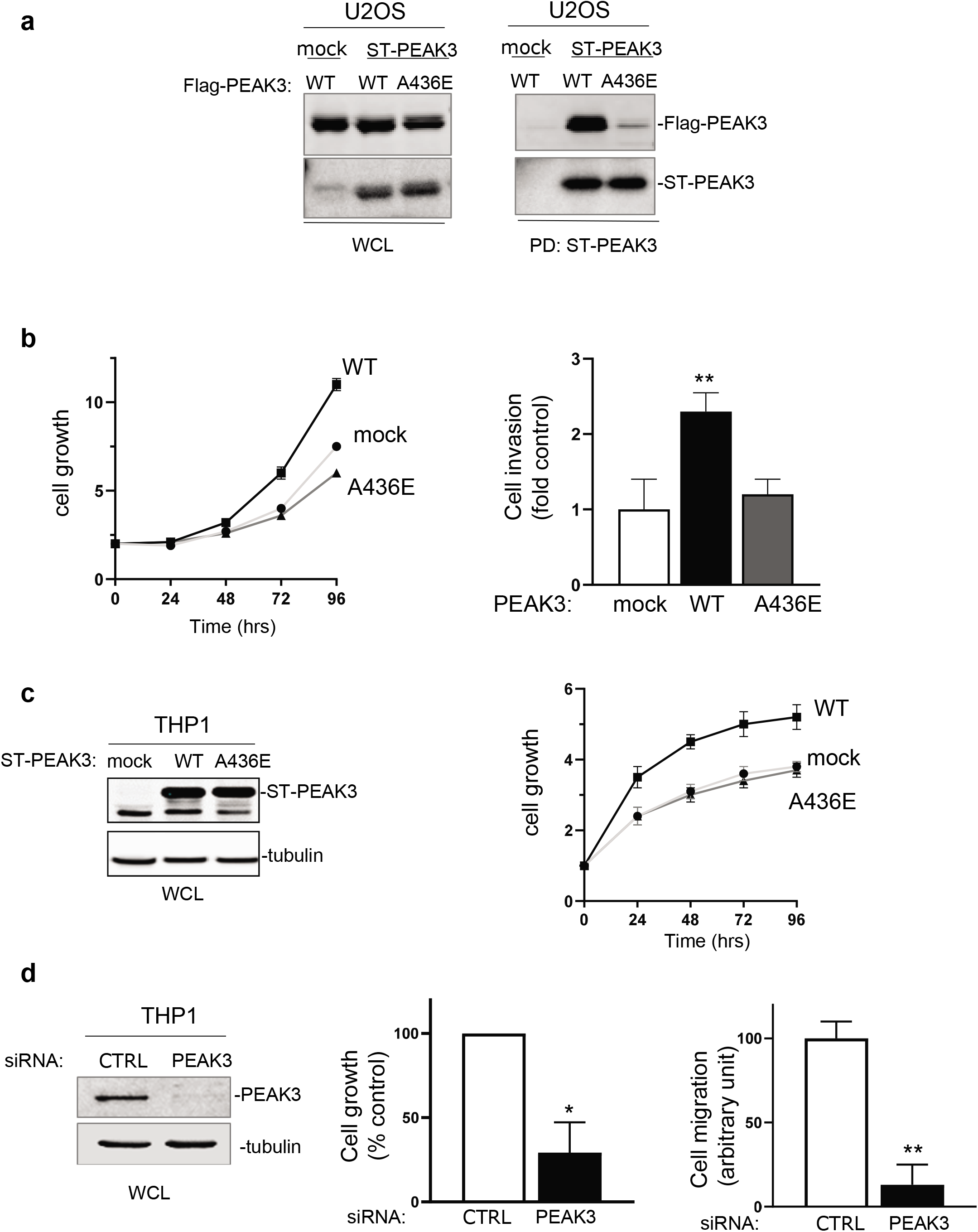
SHED-dependent PEAK3 oncogenic activity. **a.** SHED-dependent PEAK3 self-association. Co-precipitation of ST-PEAK3 on Strep-Tactin magnetic beads (PD, pull down) with FLAG-PEAK3 and A436E mutant in U2OS cells that express the indicated constructs. The level of PEAK3 in whole cell lysates (WCL) also is shown. **b.** PEAK3 expression promotes U2OS cell growth in a SHED-dependent manner. Cell growth over time (fold *vs* control) (left), and cell invasion in Boyden chambers coated with Matrigel (right) in U2OS cells that express the indicated PEAK3 constructs described in panel A (WT, wild type PEAK3; A436E, PEAK3 with an inactive SHED domain). **c**. ST-PEAK3 level (left) and cell growth over time (fold *vs* control) (left) in THP1 cells that express the indicated PEAK3 constructs. **d**. PEAK3 knockdown (left) reduces THP1 cell growth *vs* control cells (cell growth monitored for 3 days) and migration in Boyden chambers coated with an endothelial cell monolayer; *p≤0.05, **p≤0.01 (Student’s *t* test).

### PEAK3 activates AKT signalling

Next, to determine PEAK3 oncogenic signalling, we probed a phospho-kinase antibody array from THP1 cells transfected with a control siRNA or a siRNA targeting *PEAK3* (Fig. 4a). We found that PEAK3 downregulation affected the activating phosphorylation of adhesive TKs of the Src (SFK) and FAK families (Fig. S6a), and also reduced MAPK and AKT activating phosphorylations (Fig. 4a and S6a). We confirmed these results by western blotting (Fig. 4a, right panel). *PEAK3* silencing also reduced phosphorylation of the probed AKT substrates (i.e. GSK-3 alpha/beta, PRAS40 and CREB), suggesting that this pseudo-kinase is a novel upstream regulator of this pathway (Fig. S6a). Consistently, AKT activity (i.e. pS476-AKT level) was increased by about 2-fold upon PEAK3 overexpression, and reduced by 3-fold upon PEAK3 knockdown (Fig 4b). We obtained similar results in PEAK3-overexpressing U2OS cells, suggesting that PEAK3 might be a general inducer of AKT signalling (Fig. 4c). PEAK3 overexpression induced a modest increase in AKT signalling (i.e. AKT activity and AKT substrate phosphorylation). Conversely, PEAK3 strongly activated AKT in serum-starved conditions (Fig. 4c). We did not observe this effect with the PEAK3 A436E mutant, further supporting the essential role of the SHED module in PEAK3 signalling (Fig. 4c). PEAK3-AKT signalling was abrogated by pharmacological inhibition of PI3K activity (Fig. S6b), suggesting that PEAK3 is upstream to PI3K. Collectively, these data indicate that PEAK3 induces PI3K/AKT signalling, even in the absence of growth factors.

**Figure 4.**
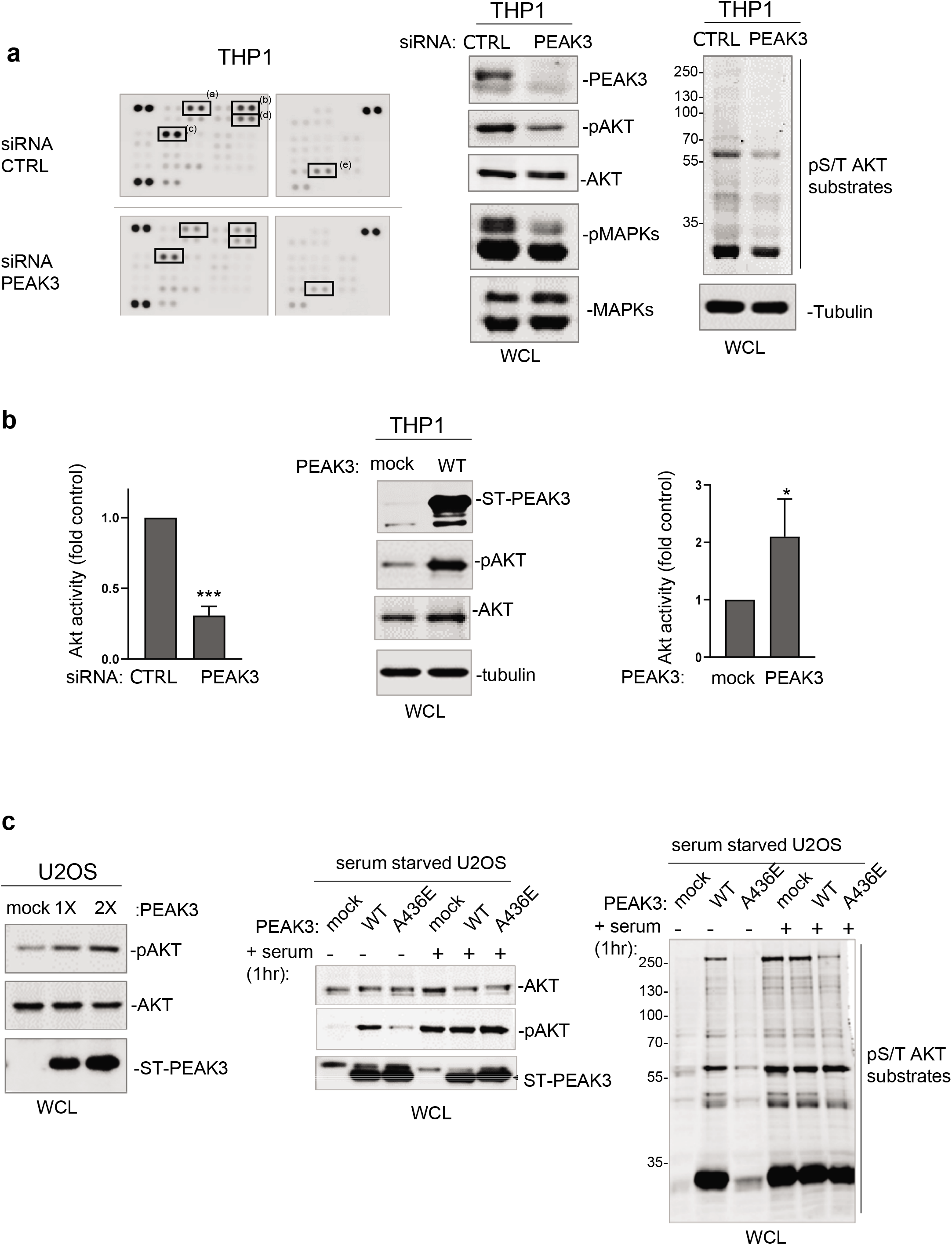
PEAK3 activates AKT signalling in the absence of growth factors. **a**. (left) Phospho-kinase array analysis of THP1 cells transfected with siRNA control (CTRL) or siRNA against PEAK3. The phospho-signal reduction in the siRNA PEAK3 condition is highlighted: (a) pT202/Y204 ERK1/2; (b) pS21/9 GSK3β; (c) pS133 CREB; (d) pS473 AKT; and (e) pT60 WNK. (right) Western blot analysis of MAPK, AKT activity and signalling (pS/T AKT substrates level). Tubulin level also is shown. **b.** (left) Quantification of AKT inhibition upon PEAK3 silencing. PEAK3 overexpression increases AKT activity in THP1 cells. A representative western blot (middle) and quantification of AKT activation (right). **c.** PEAK3 expression induces AKT signalling in U2OS cells, even in the absence of growth factors. (left) AKT activation upon transient PEAK3 expression (1X: 1μg, 2X: 2 μg of PEAK3 construct transfected). SHED-dependent PEAK3 activation of AKT signalling in the absence of growth factors. U2OS cells that stably express WT or A436E PEAK3 were serum-starved for >20 hours and stimulated or not with 10% FCS for 1 hour. Then, the levels of AKT, phosphorylated AKT (middle) and pS/T AKT substrates were assessed (right). Quantification data are the mean ± SD (n=3); *p<0.05; **p<0.01 (Student’s *t* test).

### SHED-dependent PEAK3 scaffolding activity

To further elucidate the mechanism by which PEAK3 induces AKT signalling, we performed PEAK3 interactomic analyses of U2OS, HeLa and THP1 cells that stably express ST-PEAK3 (Fig. 5a, S4b and Table S2). Label-free LC-MS/MS analysis of ST-PEAK3 complexes purified on streptavidin beads identified many signalling proteins in the three PEAK3 interactomes, including GRB2, ADP-ribosylation factor (ARF) GTPase-activating proteins ASAP1 and 2, and another member of the PEAK family (i.e. PEAK1 in the HeLa and U2OS cell interactomes, and PEAK2 in the THP1 cell interactome). This proteomic analyses identified also many 14-3-3 proteins, which regulate phospho-dependent signalling activity (25), and voltage-dependent anion-selective channel proteins (i.e. VDAC1-3), which regulate cell volume and apoptosis (Table S3) (26). Additionally, we detected the FAK-like TK PYK2 in the PEAK3 interactome of THP1 cells (Table S3). We next confirmed some of these findings by western blotting (Fig. 5b-c). Although the CRKII cytoskeletal proteins were previously reported as PEAK3 binders (22), we could not detect any endogenous CRKII protein associated with PEAK3 in our proteomic and biochemical analyses (Fig. 5c and Table S3). All these interactions required an intact SHED module (Fig. 5c), suggesting that PEAK3 dimerization is essential for interaction with these signalling proteins. As ASAP1/2, VDACs and PYK2 have not been detected in the previously reported PEAK1 and PEAK2 interactomes (9,21), PEAK1-3 may recruit a unique set of signalling proteins to induce intracellular signalling.

**Figure 5.**
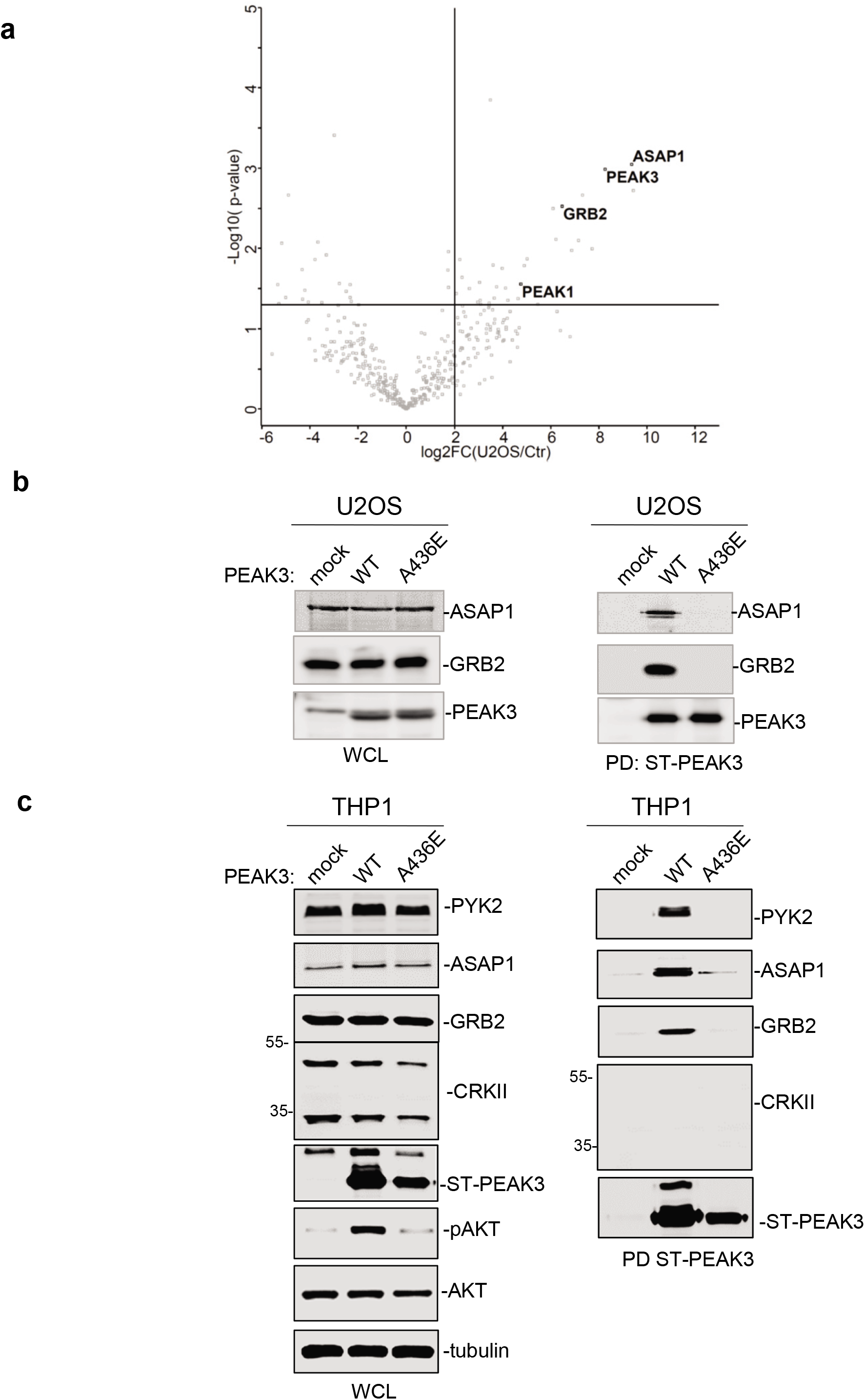
PEAK3 interactomics identified GRB2, ASAP1/2 and PYK2 as SHED-dependent binders. **a.** Interactome of PEAK3 in U2OS cells represented as volcano plots of the MS data. **b** and **c.** Validation by western blot analysis of PEAK3 interaction with the indicated signalling proteins in U2OS (b) and THP1 cells (c).

### PEAK3 activates PYK2 to promote AKT signalling

The fact that PEAK3 interacts with PYK2 suggests that like PEAK2 (5,9), PEAK3 may activate specific TKs to mediate phospho-tyrosine signalling. We tested this hypothesis by co-expressing PEAK3 and PYK2 in HEK293T cells. While PEAK3 alone did not clearly affect protein tyrosine phosphorylation, PEAK3 robustly activated ectopic PYK2, as measured on the phosphorylation level of regulatory tyrosine 402 and 881 (Fig 6a) (27). Interestingly, PYK2 activation was accompanied by an increase in cellular protein tyrosine phosphorylation, including a 50 KD and 120KDa band (Fig 6a). PEAK3 complex affinity-purification next identified PEAK3 and PYK2 as ones of these tyrosine phosphorylated proteins (Fig 6b). This result indicates that PEAK3 can undergo tyrosine phosphorylation and suggests that PYK2 acts as an upstream TK. This PEAK3 activating mechanism on PYK2 was next confirmed in PEAK3 overexpressing U2OS cells (Fig 7a). Importantly, this molecular effect was further enhanced in serum starved conditions (Fig 7a), supporting also a growth factor-independent mechanism of PEAK3 activation of PYK2. PEAK3 knockdown reduced PYK2 activity in THP1 cells (Fig 7b), supporting the existence of a similar endogenous PEAK3-PYK2 signalling in leukemic cells. Having demonstrated the existence of PEAK3-AKT and PEAK3-PYK2 signalling, we then asked whether PYK2 would mediate the growth-independent AKT activation by PEAK3. Acute pharmacological inhibition of PYK2 (PYK2i) in U2OS reduced by 3-fold the stimulating effect of PEAK3 on AKT, while this inhibitor had no effect on AKT activity in control cells (Fig 7c). Overall, these findings demonstrate that PEAK3 activates PYK2 and support a role of PYK2 activity in PEAK3-AKT signalling.

**Figure 6.**
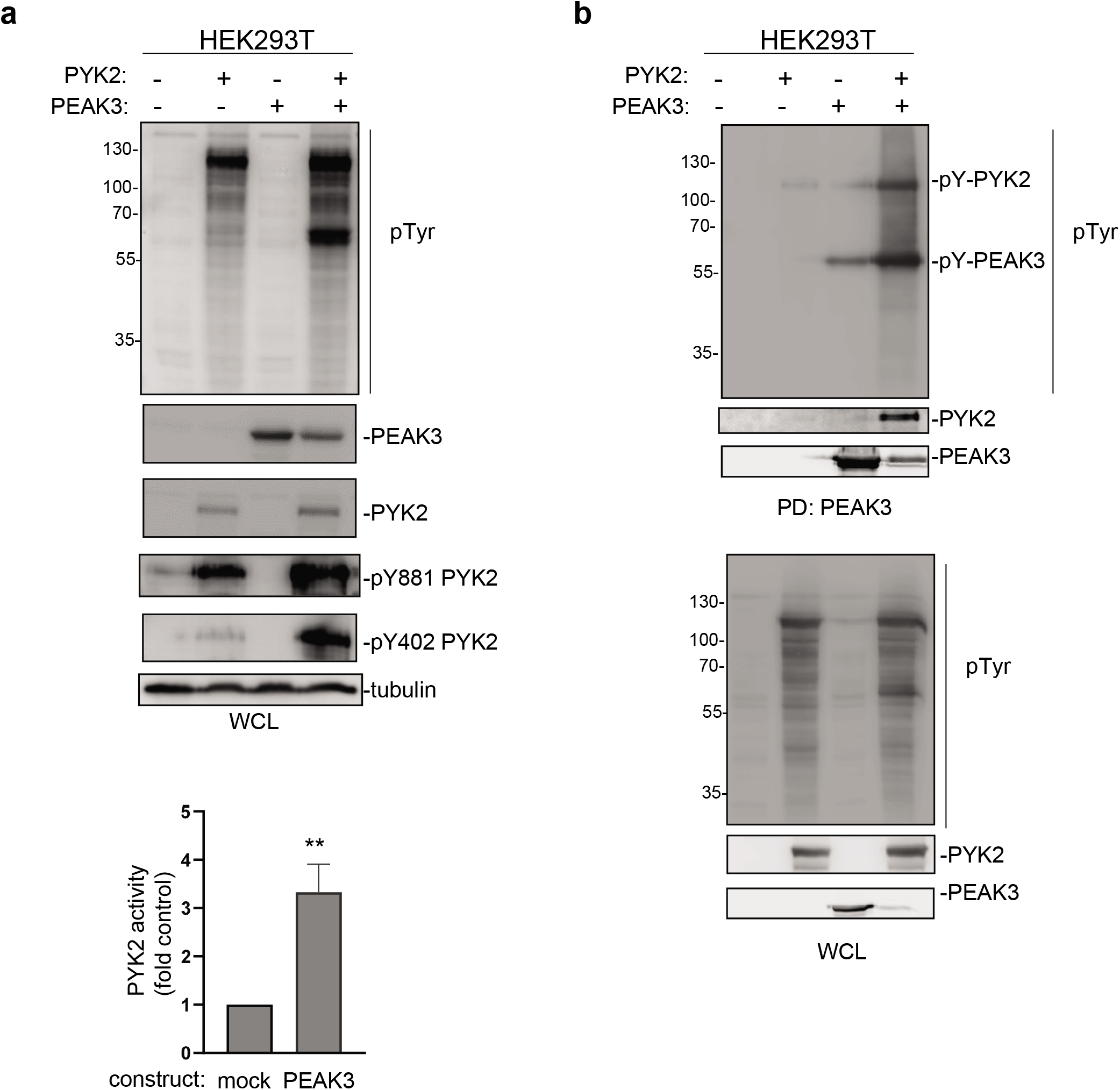
PEAK3 activates PYK2. **a.** PYK2 activation by PEAK3 in HEK293T cells. The level of protein tyrosine phosphorylation, PEAK3 and PYK2 and phosphorylated PYK2 (pPYK2; pTyr402 and pTyr881) was assessed in cells transfected with the indicated constructs for 40 hours. (left) Representative example; (right) Quantification of three independent experiments. **p<0.01 (Student’s *t* test). **b**. PEAK3 tyrosine phosphorylation by PYK2. The level of PEAK3 and PYK2 protein tyrosine phosphorylation was assessed in ST-PEAK3 complexes purified on streptavidin beads from cells transfected with indicated constructs for 40 hours.

**Figure 7.**
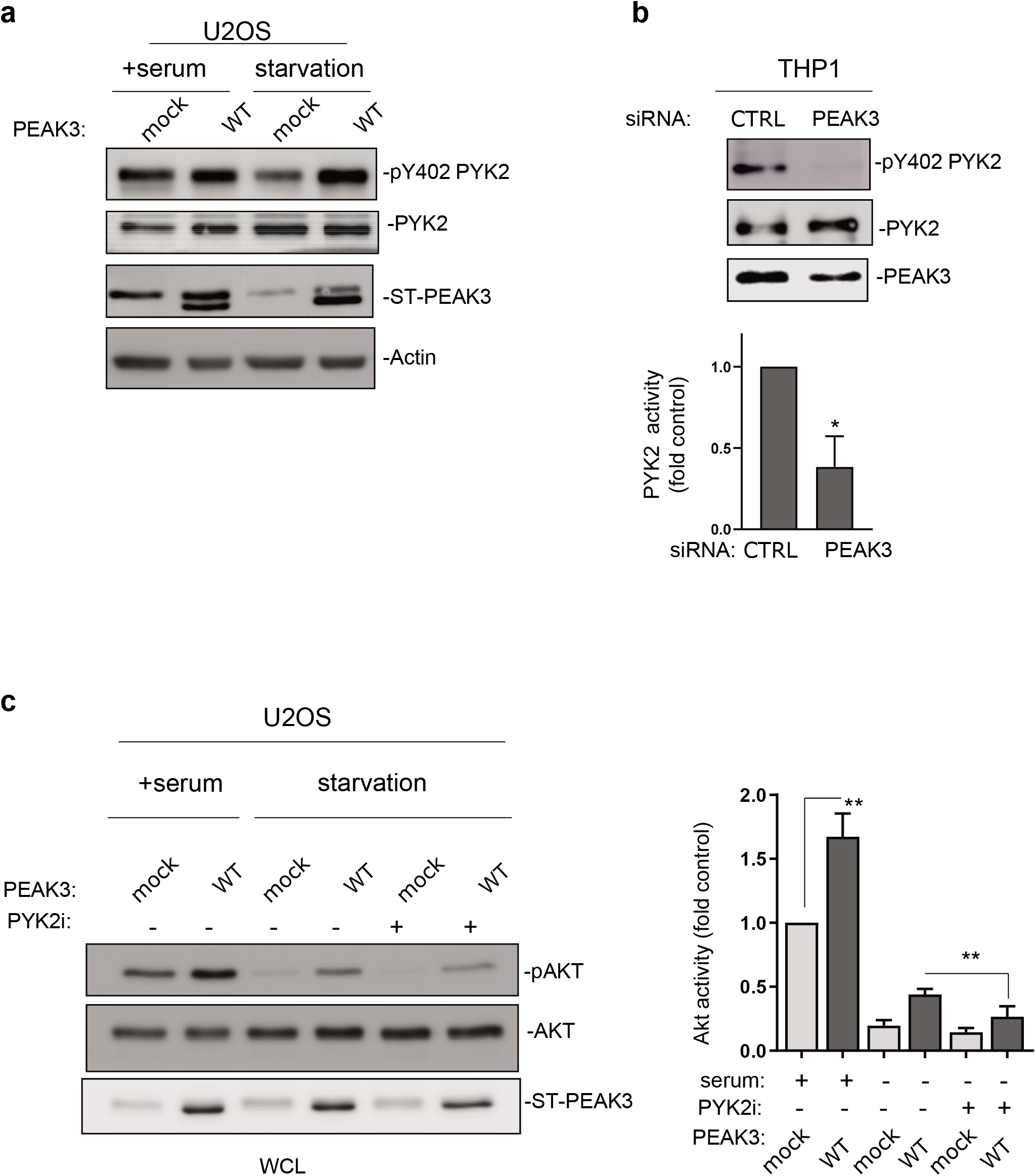
PYK2 activity mediates PEAK3-AKT signalling. **a.** PYK2 activation by PEAK3 stably overexpressed in U2OS cells that were serum-starved or not for >20 hours. The level of PEAK3, PYK2 and pPYK2 was assessed by western blotting. **b.** Regulation of PYK2 activity by endogenous PEAK3 in THP1 cells. PYK2 activity (pTyr402 PYK2 level) was measured in cells transfected with the indicated siRNAs. (bottom) Quantification of PYK2 activity (mean ± SD of three independent experiments); **p<0.01 (Student’s *t* test). **c**. PYK2 inhibition reduces PEAK3 activation of AKT in the absence of growth factors. U2OS cells that express or not PEAK3 (were serum-starved overnight or not, and incubated with the PYK2 inhibitor (PYK2i) PF-431396 (1μM) or vehicle (DMSO). The level of AKT, PEAK3 and phosphorylated AKT was then quantified (mean ± SD of three independent experiments); **p<0.01 (Student’s *t* test) (left).

## DISCUSSION

Here we showed that the pseudo-kinase PEAK3 has an oncogenic function, as previously reported for PEAK1 and 2. PEAK3 overexpression enhanced cancer cell growth and migration, while its silencing reduced these pro-tumour effects. PEAK3 expression profile suggested a restricted function in myeloid cells, such as granulocytes, and our results revealed a pro-tumoral function in AML. This finding was supported by the aberrant PEAK3 expression observed in AML patient samples and by the finding that endogenous PEAK3 is implicated in the regulation of THP1 leukemic cell features. As PEAK1/2 can mediate TK signalling induced by growth factor receptors (5,6), the absence of association between PEAK3 and the oncogenic receptor TK FLT3-TD in AML (28) is surprising. This suggests that PEAK3 may not be a critical component of receptor TK signalling or that it displays redundant signalling functions. This hypothesis is corroborated by the finding that PEAK3 activates AKT signalling in the absence of growth factors. Moreover, the heterogeneous PEAK3 expression in AML patient samples suggest a PEAK3 function in a specific AML subtype that needs to be thoroughly characterized. The association between PEAK3 expression and DNTM3A (28) is consistent with this hypothesis.

This study also uncovers an essential role of the SHED module in PEAK3 biological activities. Specifically, PEAK3 dimerization was necessary for PEAK3 pro-tumoral activity, intracellular signalling, and localization to FAs. PEAK3 dimerization was also essential for interaction with important signalling proteins, uncovering a general scaffolding mechanism regulated by this dimeric domain to allow protein interactions. As this domain is conserved among PEAK family member, we might predict a similar function in PEAK1 and PEAK2 signalling that needs to be confirmed. Daly et al reported a hetero-dimerization mechanism involved in PEAK1/2 signalling specificity (21). We did not address here whether this mechanism contributes also to PEAK3 signalling. Nevertheless, the presence of another PEAK member in our PEAK3 interactomes suggests that heterodimers may contribute to PEAK3 signalling. However, as PEAK3 was the main PEAK member expressed in blood cells (Fig. S1), PEAK3 signalling may primarily occur via PEAK3 homo-dimerization. Our imaging analysis revealed that nuclear PEAK3 localization is not affected by protein dimerization. This suggests that the SHED module does not regulate all PEAK3 functions and that PEAK3 has an additional nuclear function, as described for PEAK2 (29), that remains to be characterized.

One important finding from our molecular study is the identification of PEAK3 as a novel upstream activator of AKT signalling, which plays an essential role in AML development (30). This result corroborates a previous report showing that PEAK3 is a potential regulator of AKT signalling using a shRNA screening approach (31), although this observation was not further investigated. Our results also suggest that PEAK3 might define an important mechanism of AKT activation in the absence of growth factors. This mechanism could be essential for malignant cell survival and dissemination during disease progression. Our interactomic analysis also showed that PYK2 is an important mediator of this signalling cascade. This is consistent with FAK-like functions described in AML, where high FAK expression level is associated with enhanced blast migration, increased cellularity, and poor prognosis (32). How PYK2 mediates PEAK3-AKT signalling is unknown. However, the fact that ASAP1 is a new PEAK3 binder and a PYK2 substrate that can activate AKT signalling suggests that PYK2 promotes AKT activation via ASAP1 (33–35). Therefore, PEAK3 might act as a scaffold to bring PYK2 and ASAP1 together for efficient ASAP1 phosphorylation and AKT signalling. Moreover, our results suggest that PEAK3 activates PYK2 through a SHED-dependent mechanism, as previously reported for PEAK2 activation of CSK (9). Therefore, we propose that, in addition to its scaffolding function, PEAK3 and PEAK2 use an allosteric mechanism to activate associated TKs for signalling. One important question not addressed in our study is the mechanism of PEAK3 regulation. During the preparation of this manuscript, Daly et al reported a similar PEAK3 oncogenic function in breast epithelial cells and showed that PEAK3 tyrosine phosphorylation promotes GBR2 and ASAP1 binding (36). Although many upstream TKs, including SFKs, could phosphorylate PEAK3, our finding suggest that PYK2 phosphorylates PEAK3, like CSK phosphorylates PEAK2. The precise mechanism by which PEAK3 activates PYK2 deserves further investigation; however, as PYK2 is regulated by dimerization (27), its binding to PEAK3 might facilitate this activation mechanism.

Finally, our study brings a unifying model on how PEAK pseudo-kinases regulate oncogenic signalling. GRB2 and possibly CRK proteins are common PEAK1-3 binders (9,21,22), through a conserved proline-rich sequence at their N-terminus. Moreover, our interactomic analyses identified a unique set of signalling proteins recruited by PEAK3 that were not detected in previous PEAK1/2 interactomes (9,21). Altogether, these findings support the existence of a common SHED-dependent mechanism of PEAK1-3 oncogenic activity mediated through the recruitment of a unique set of signalling proteins to induce specific oncogenic functions. Lopez et al found that the conserved DFG motif in the PEAK3 pseudo-kinase domain is implicated in PEAK3 dimerization (22). This raises the interesting idea that besides the SHED module, the pseudo-kinase domain also might participate in PEAK3 signalling. Consistent with this idea, recent molecular studies showed that despite the inaccessible ATP site found in crystallography, pseudo-kinases display higher kinase flexibility than previously expected, to enable signalling (2,4,37). Therefore, small molecules that affect pseudo-kinase dynamics may inhibit their function (4,37). Whether PEAK1-3 signalling is regulated by a similar mechanism deserves investigation. In this case, these pseudo-kinases could represent attractive therapeutic targets in human cancers, including leukaemia.

## Authors’ contribution

All authors contributed extensively to the work presented in this paper. Experimental analysis and data acquisition: YO, EF and YB. Proteomic and bioinformatics analyses: SU (proteomic), NG (transcriptomic) and PF (phylogenetic). PEAK3 analyses in AML: ES, NG and JES. CR provided human samples. Project supervision: SR and DF. Writing the paper: SR.

## Acknowledgment

This work was supported by La Ligue Contre le Cancer (Equipe Labellisée LIGUE2017 and 2020), CNRS, and the University of Montpellier. SR is an INSERM investigator. We thank Raphael Gaudin with his help with the cell transmigration assays, the MRI and FPP facilities of Montpellier Biocampus (www.biocampus.cnrs.fr) for imaging and proteomic analyses respectively, Gilles Labesse (CBS, Montpellier) and our colleagues for helpful discussion.

