## supplementary information for "SHED-dependent oncogenic signalling of the PEAK3 pseudo-kinase"

**Tables S1. Accession numbers of the PEAK pseudo-kinases used in this study.**

**Tables S2. Characteristics of AML patient samples.**

**Table S3. Main hits recovered from the PEAK3 interactomic analyses in U2OS, HeLa and THP1 cells.**

**Figure S1. *PEAK1-3* mRNA distribution in human tissues.** Normalized *PEAK1-3* expression levels from the Human Protein Atlas (24).

**Figure S2. Characterization of our anti-PEAK3 antibody a.** Western blot (WB) analysis with our anti-PEAK3 antibody of a total lysate of U2OS cells that overexpress ST-PEAK3 and that was transfected with indicated siRNA (left). Immunoprecipitation (ip) of ST-PEAK3 overexpressed in HeLa cells with our anti-PEAK3 antibody (right); WCL, whole cell lysate; FT, Flow Through. **b.** The anti-PEAK3 antibody recognizes endogenous PEAK3 in a THP1 cell lysate. The antibody specificity is shown using THP1 cells transfected with a siRNA targeting PEAK3. **c.** Western blot analysis with our anti-PEAK3 antibody of PEAK3 expression in whole cell lysates of the indicated cell lines. The antibody specificity is shown using THP1 cells transfected with a siRNA targeting PEAK3. Both low and longer exposures are shown.

**Figure S3. PEAK3 transcript level in AML patient samples. a.** FAB morphology in BEAT AML. **b.** Association between PEAK3 aberrant expression and indicated specific oncogenic mutations.

**Figure S4. PEAK3 signalling in HeLa cells. a**. SHED-dependent PEAK3 self-association. Co-precipitation of ST-PEAK3 (wild type, WT, and the indicated mutant) on streptavidin beads (PD, pull down) with FLAG-PEAK3 from HeLa cells that express the indicated constructs. PEAK3 expression in whole cell lysates (WCL) is also shown. **b.** PEAK3 interactome in HeLa cells represented as volcano plots of the MS data.

**Figure S5. PEAK3 cellular distribution. a.** Direct fluorescence of U2OS cells that express GFP-PEAK3 (WT or A436E mutant). Merge, co-localization of GFP-PEAK3 with paxillin. **b**. PEAK3 localization by indirect anti-HA immunofluorescence in U2OS cells that express HA-ST-PEAK3 (wild type or the A436E mutant).

**Figure S6. PEAK3-AKT signalling. a**. Quantification of the selected signals from the PEAK3-dependent phospho-kinase array in THP1 cells. **b.** PEAK3 activation of AKT in U2OS cells is inhibited by incubation with the pan-PI3K inhibitor LY294002.

**Figure S1**

### PEAK1

**RNA tissue specificity:** Low tissue specificity

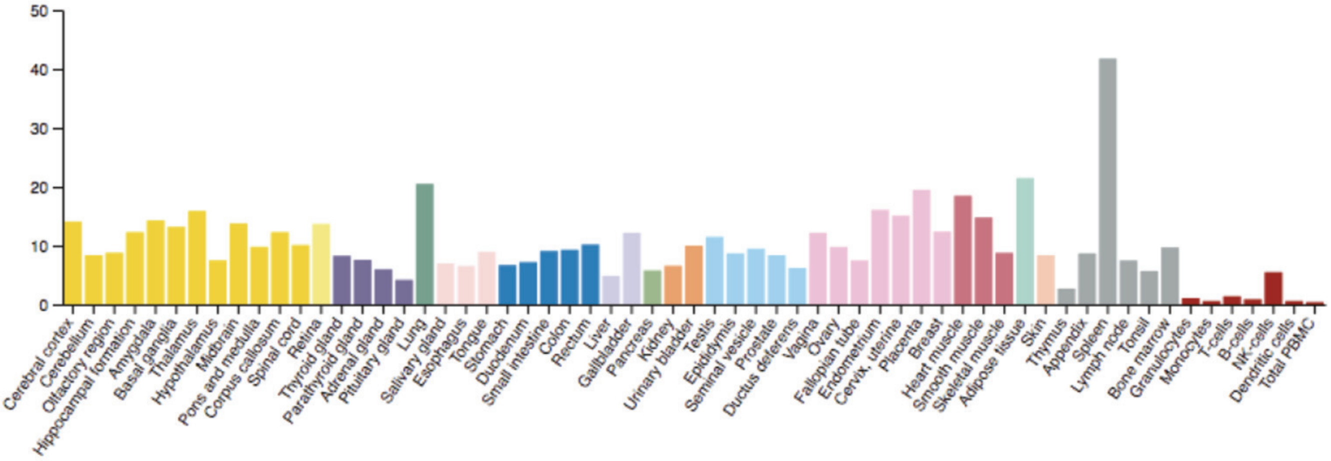


### PEAK2

**RNA tissue specificity:** Tissue enhanced (brain)

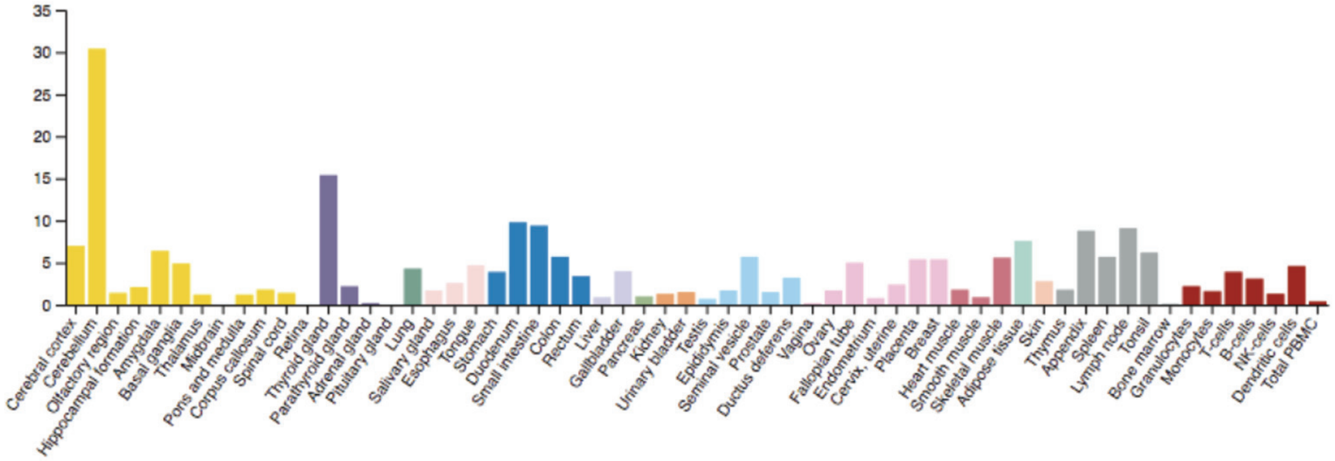


### PEAK3

**RNA tissue specificity:** Tissue enhanced (blood, lymphoid tissue)

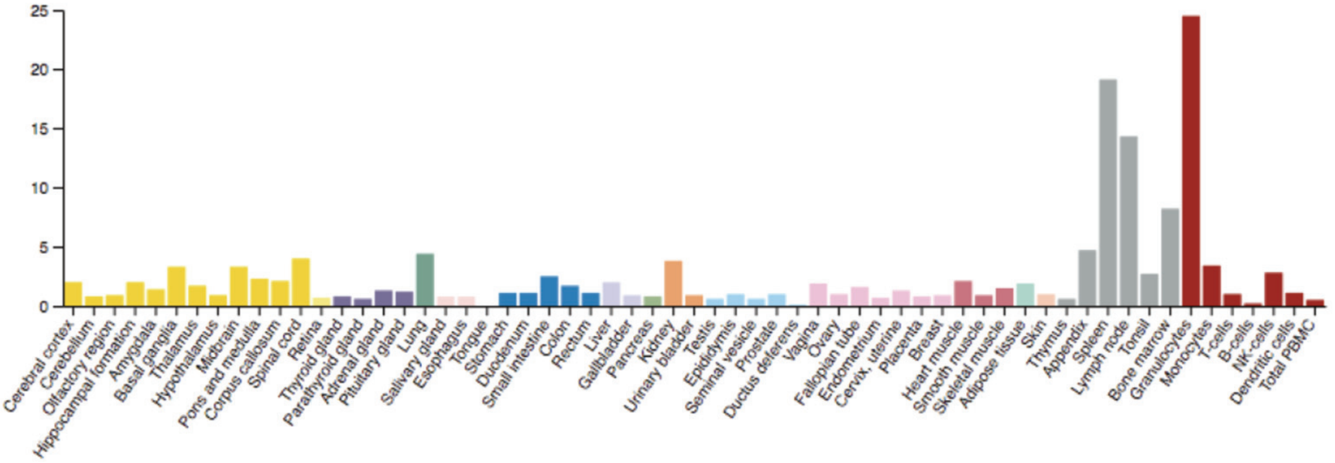

**Figure S2**

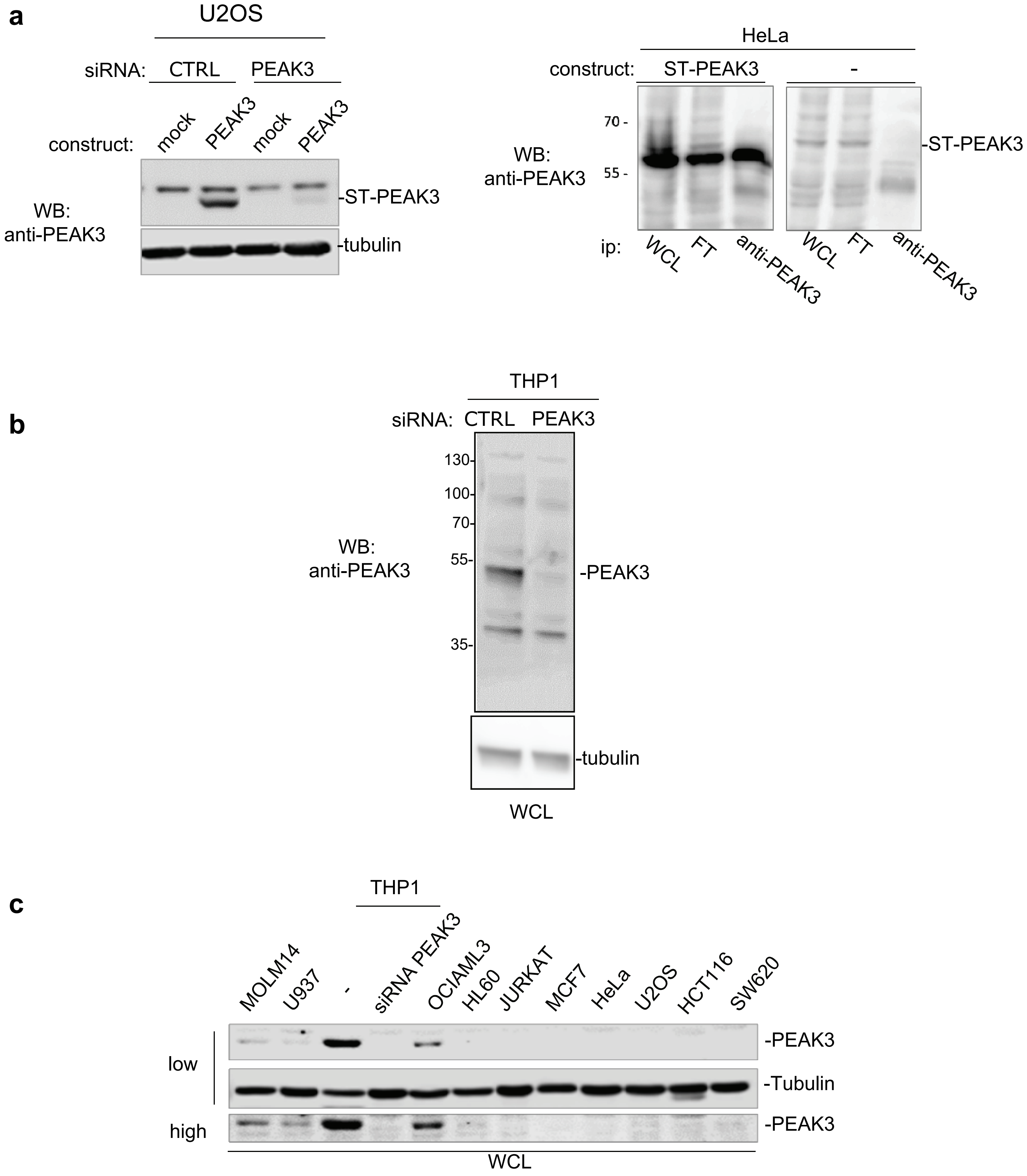
a
U2OS
siRNA: CTRL PEAK3
construct: mock PEAK3 mock PEAK3
WB: anti-PEAK3
-ST-PEAK3
-tubulin
HeLa
construct: ST-PEAK3 -
70 -
55 -
WB: anti-PEAK3
-ST-PEAK3
ip: WCL FT anti-PEAK3 WCL FT anti-PEAK3
b
THP1
siRNA: CTRL PEAK3
130-
100-
70-
55-
35-
WB: anti-PEAK3
-PEAK3
-tubulin
WCL
c
THP1
MOLM14
U937
-
siRNA PEAK3
OCIAML3
HL60
JURKAT
MCF7
HeLa
U2OS
HCT116
SW620
low
high
-PEAK3
-Tubulin
-PEAK3
WCL

**Figure S3**

**a**

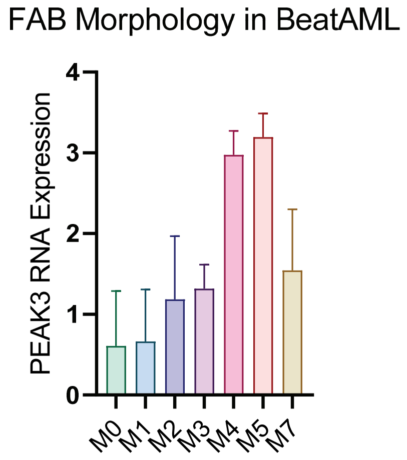
FAB Morphology in BeatAML
PEAK3 RNA Expression
0
1
2
3
4
M0
M1
M2
M3
M4
M5
M7

**b**

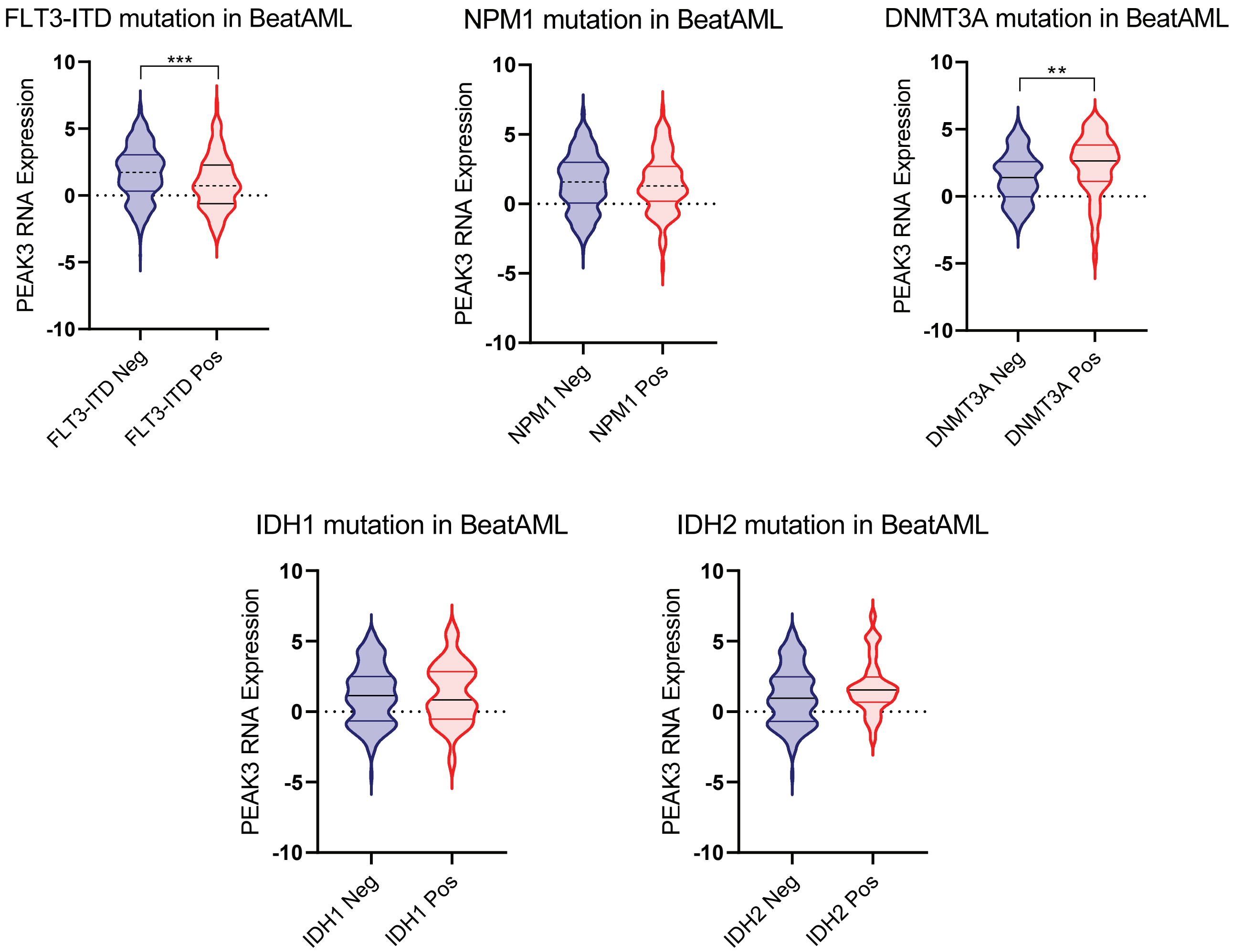
FLT3-ITD mutation in BeatAML
PEAK3 RNA Expression
***
FLT3-ITD Neg
FLT3-ITD Pos
NPM1 mutation in BeatAML
NPM1 Neg
NPM1 Pos
DNMT3A mutation in BeatAML
**
DNMT3A Neg
DNMT3A Pos
IDH1 mutation in BeatAML
IDH1 Neg
IDH1 Pos
IDH2 mutation in BeatAML
IDH2 Neg
IDH2 Pos
10
5
0
-5
-10

**Figure S4**

a

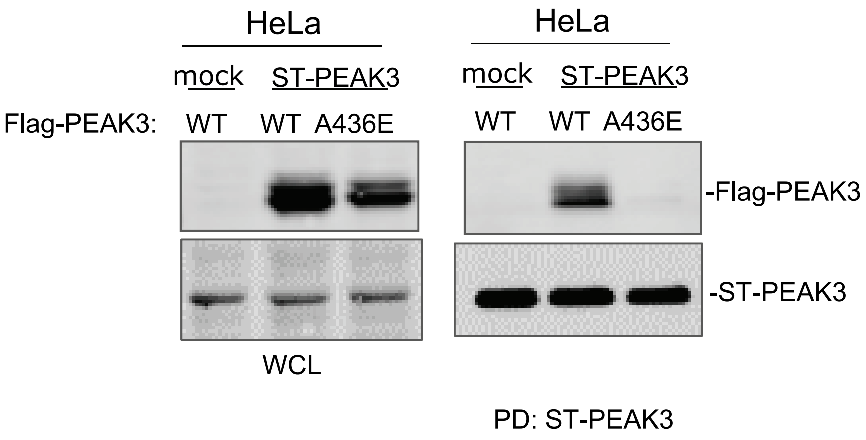
HeLa
mock
ST-PEAK3
HeLa
mock
ST-PEAK3
Flag-PEAK3:
WT
WT
A436E
WT
WT
A436E
-Flag-PEAK3
-ST-PEAK3
WCL
PD: ST-PEAK3

b

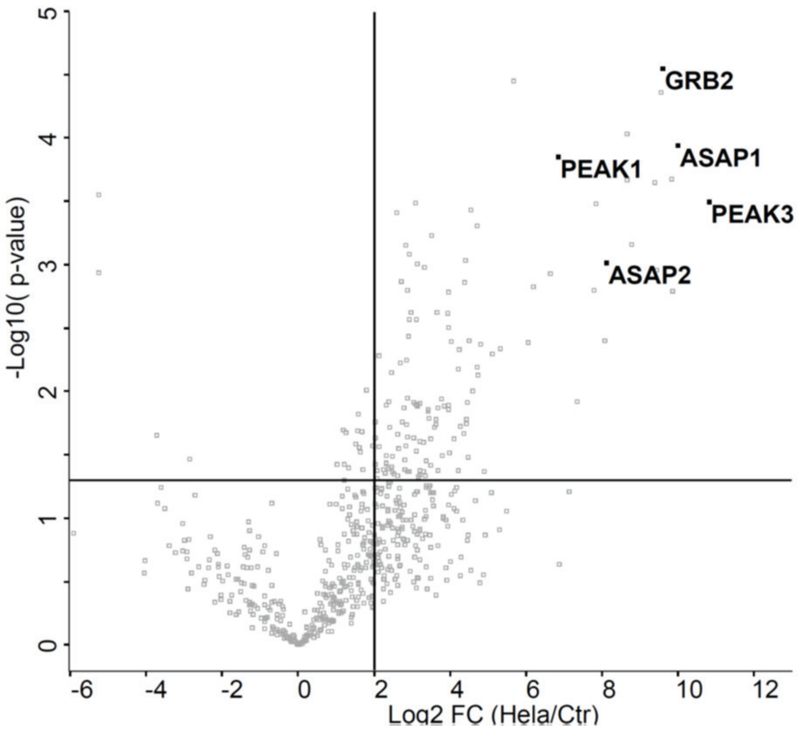
GRB2
PEAK1
ASAP1
PEAK3
ASAP2
-Log10( p-value)
Log2 FC (Hela/Ctr)

**Figure S5**

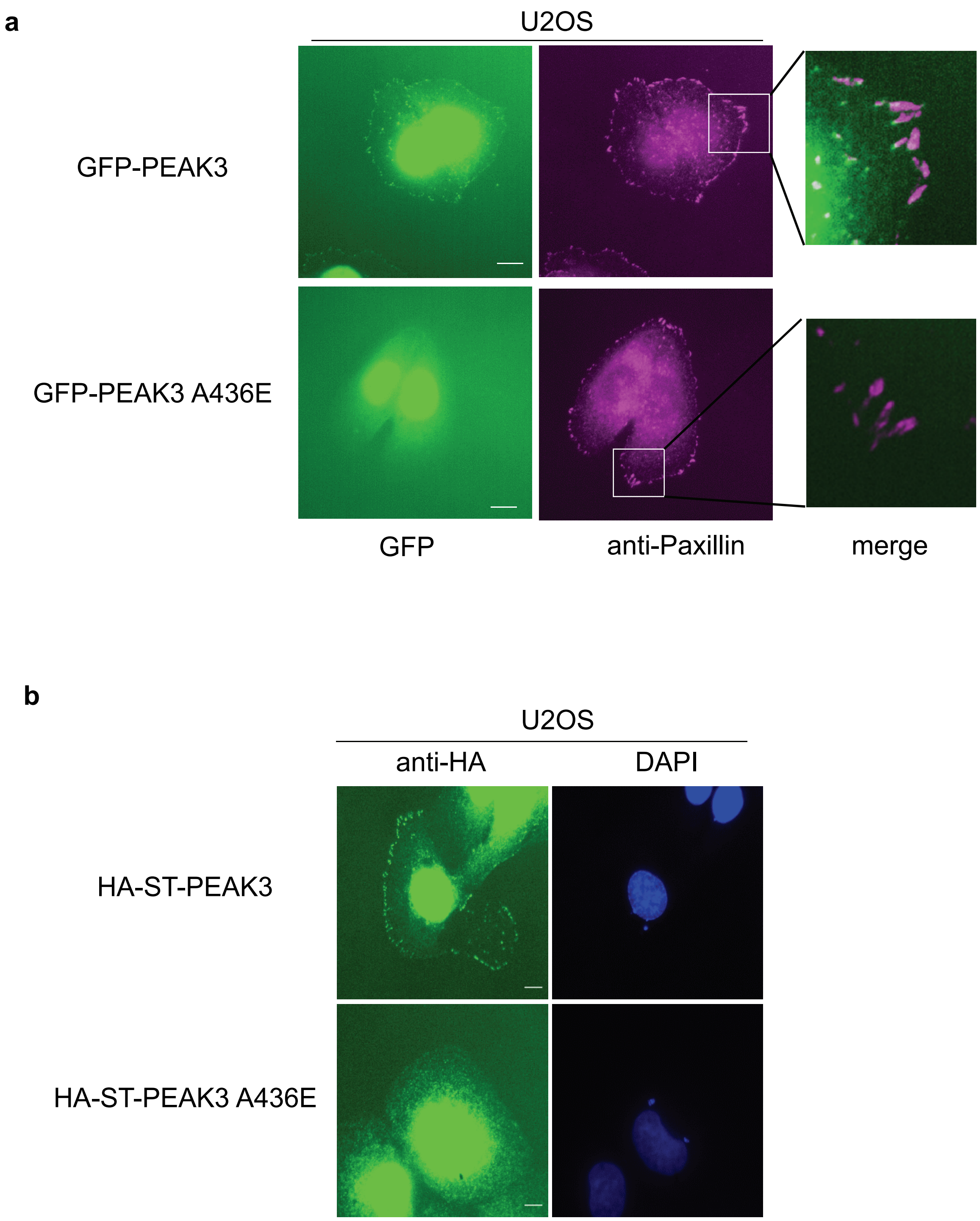
a
U2OS
GFP-PEAK3
GFP-PEAK3 A436E
GFP
anti-Paxillin
merge
b
U2OS
anti-HA
DAPI
HA-ST-PEAK3
HA-ST-PEAK3 A436E

**Figure S6**

**a**

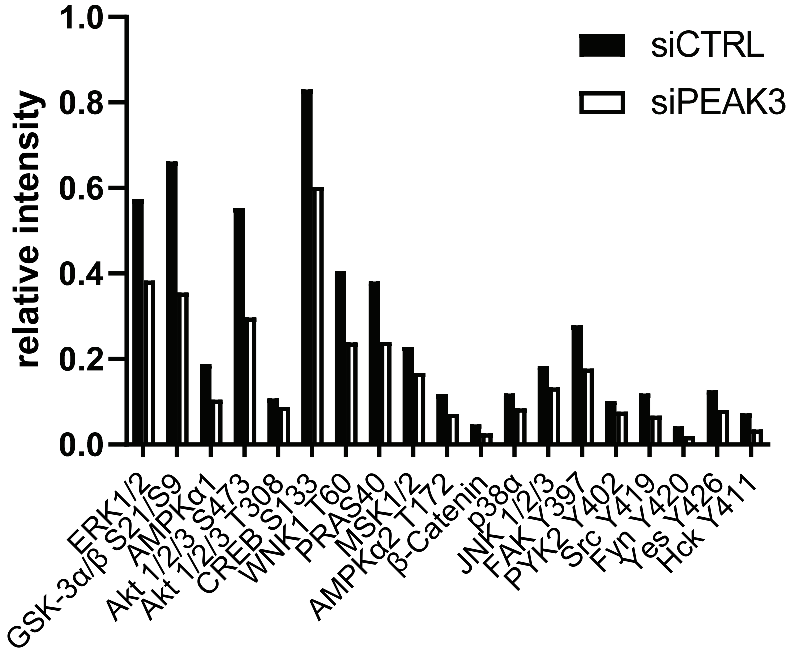
relative intensity
1.0
0.8
0.6
0.4
0.2
0.0
siCTRL
siPEAK3
ERK1/2
GSK-3α/β S21/S9
AMPKα1
Akt 1/2/3 S473
Akt 1/2/3 T308
CREB S133
WNK1 T60
PRAS40
MSK1/2
AMPKα2 T172
β-Catenin
p38α
JNK 1/2/3
FAK Y397
PYK2 Y402
Src Y419
Fyn Y420
Yes Y426
Hck Y411

**b**

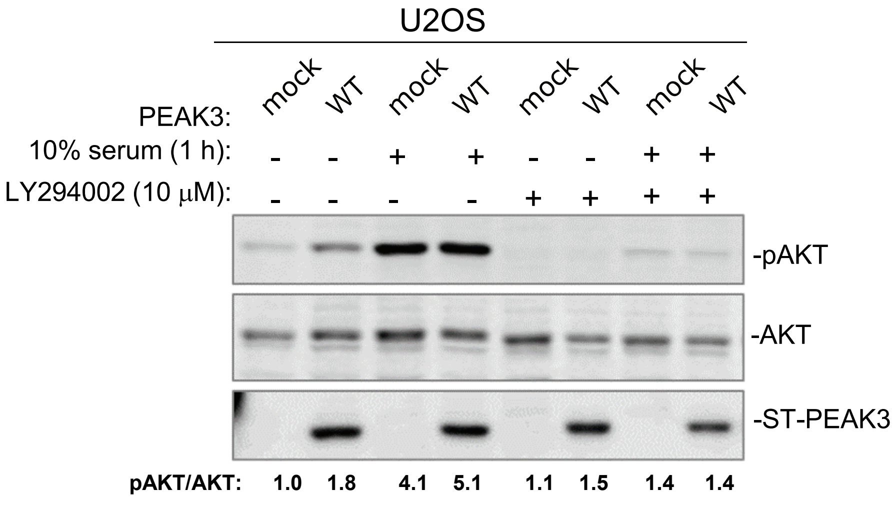
U2OS
PEAK3: mock WT mock WT mock WT mock WT
10% serum (1 h): - - + + - - + +
LY294002 (10 µM): - - - - + + + +
-pAKT
-AKT
-ST-PEAK3
pAKT/AKT: 1.0 1.8 4.1 5.1 1.1 1.5 1.4 1.4
